# Characterisation of the HIV-1 Molecular Epidemiology in Nigeria: Origin, Diversity, Demography and Geographic Spread

**DOI:** 10.1101/410431

**Authors:** Jamirah Nazziwa, Nuno Faria, Beth Chaplin, Holly Rawizza, Patrick Dakum, Alash’le Abimiku, Man Charurat, Nicaise Ndembi, Joakim Esbjörnsson

## Abstract

Nigeria has been reported to have the highest number of AIDS-related deaths in the world. In this study, we aimed to determine the HIV-1 genetic diversity and phylodynamics in Nigeria. We analysed 1442 HIV-1 *pol* sequences collected 1999-2014 from four geopolitical zones in Nigeria. Phylogenetic analysis showed that the main circulating strains was the circulating recombinant strain (CRF) 02_AG (44% of the analysed sequences), subtype G (8%), and CRF43_02G (16%); and that these were introduced in Nigeria in the 1960s, 1970s and 1980s, respectively. The number of effective infections decreased in Nigeria after the introduction of free antiretroviral treatment in 2006. We also found a significant number of unique recombinant forms (22.7%). The majority of those were recombinants between two or three of the main circulating strains. Seven of those recombinants may represent novel CRFs. Finally, phylogeographic analysis suggested multiple occasions of HIV-1 transmissions between Lagos and Abuja (two of the main cities in Nigeria), that HIV-1 epidemic started in these cities, and then dispersed into rural areas.

**IMPORTANCE:** Nigeria has the second largest HIV-1 epidemic in the world with the highest number of AIDS-related deaths. The few previous reports have focused on local HIV-1 subtype/CRF distributions in different Nigerian regions, and the molecular epidemiology of HIV-1 in Nigeria as a whole is less well characterized. In this study, we describe the HIV-1 spatiotemporal dynamics of the five dominating transmission clusters representing the main characteristics of the epidemiology. Our results may contribute to inform prevention strategies against further spread of HIV-1 in Nigeria.

## INTRODUCTION

Thirty-seven years after the first AIDS cases were described^1,2^, HIV-1 is still a major public health problem that affects approximately 36.7 (30.8–42.9) million people globally^3^. Sub-Saharan Africa accounts for approximately 70% of all those infections. Nigeria, the most populous country in this region, has been ranked as the country with the highest number of AIDS-related deaths and the second highest number of HIV-1 infected cases in the world^4,5^. HIV-1 serological surveys in Nigeria were initiated in 1991, and an adult prevalence of 1.8% (760,000) was reported^6^. This figure gradually increased to 5.8% in 2001 (2.6 million), before declining to 2.9% in 2016 (3.1 million)^3,6^. Previous reports on circulating HIV-1 strains in Nigeria have identified subtype G and the circulating recombinant form (CRF) 02_AG as the most common^7-9^. In addition, CRF43_02G was recently shown to be highly prevalent in the capital Abuja^10^. However, estimates on the contribution of each strain to the Nigerian HIV-1 epidemic have varied considerably, with estimates of frequency varying between 22% and 50% for subtype G, and between 19% and 60% for CRF02_AG^8-15^. These variations may be due to differences between geographic areas and transmission groups. A clearer picture of the HIV-1 subtype/CRF distributions in Nigeria is therefore warranted. In addition, it has been suggested that the genetic composition of the infecting HIV-1 strain may influence disease progression rate, probability and efficiency of transmission, interaction with the host, response to viral treatment and vaccine development^16-23^.

Advances in molecular biology techniques, databases, bioinformatics tools and expanded disease surveillance programs have provided an opportunity for scientists to conduct HIV-1 epidemiological studies to understand the diversity, origin and transmission dynamics of HIV-1 by phylodynamic approaches^24^. Applying established epidemiological models on HIV-1 gene sequences in combination with data on time and location of sampling, epidemic growth rate, number of effective infections, timing, origin and dispersal of different HIV-1 lineages can be obtained^24-26^.

The objective of the current study was to characterize the molecular epidemiology of HIV-1 in Nigeria using a large dataset of *pol* sequences collected from 1999-2014. Phylodynamic approaches were employed to determine the HIV-1 diversity and to uncover the demographic history and the HIV-1 dissemination routes of the main circulating strains within Nigeria. This study increase the understanding of the Nigerian HIV-1 epidemic and may inform HIV-1 intervention strategies aiming at reducing the spread of HIV-1 in Nigeria.

## MATERIALS AND METHODS

### Sequence dataset

We analyzed 366 previously unpublished HIV-1 *pol* sequences (positions in HXB2 K03455: 2253-3364) collected in Abuja, Nigeria from 2006-2011 together with all Nigerian *pol* sequences from the corresponding genetic region available in the Los Alamos HIV-1 sequence database (N=1076, April 2015, http://www.hiv.lanl.gov/, Table 1). Missing information on date and location of sampling was obtained through contact with relevant research centers.

**Table 1.** Proportion of subtype/CRF in the Nigerian dataset of previously published and new sequences collected in the period 1999-2013.

| Subtype/CRF | N | % | Geography |  |  |  |  |  |  |  |  |  |  |  |  |  |  |  |  |  |  |
| --- | --- | --- | --- | --- | --- | --- | --- | --- | --- | --- | --- | --- | --- | --- | --- | --- | --- | --- | --- | --- | --- |
|  |  |  | Southwest |  |  |  |  |  |  | North Central |  |  |  |  | North East & North West |  |  |  |  |  |  |
|  |  |  | LAG | IBA | OND | OSU | OYO | ENU | TOTAL | ABV | JOS | NIG | NC | TOTAL | KAD | YOB | MAI | KAN | ADA | BOR | TOTAL |
| **A1** | 24 | 1.7 | 9 | 2 | 0 | 0 | 0 | 0 | 11 | 7 | 5 | 0 | 0 | 12 | 0 | 0 | 1 | 0 | 0 | 0 | 1 |
| **B** | 4 | 0.3 | 1 | 1 | 0 | 0 | 0 | 0 | 2 | 2 | 0 | 0 | 0 | 2 | 0 | 0 | 0 | 0 | 0 | 0 | 0 |
| **C** | 14 | 1.0 | 2 | 1 | 0 | 0 | 2 | 0 | 5 | 6 | 2 | 0 | 1 | 9 | 0 | 0 | 0 | 0 | 0 | 0 | 0 |
| **CRF02_AG** | 636 | 44.1 | 176 | 27 | 0 | 1 | 18 | 0 | 222 | 284 | 94 | 0 | 10 | 388 | 13 | 0 | 11 | 2 | 0 | 0 | 26 |
| **CRF06_cpx** | 64 | 4.4 | 18 | 2 | 0 | 0 | 0 | 0 | 20 | 36 | 6 | 0 | 1 | 43 | 0 | 0 | 1 | 0 | 0 | 0 | 1 |
| **CRF09_cpx** | 1 | 0.1 | 0 | 1 | 0 | 0 | 0 | 0 | 1 | 0 | 0 | 0 | 0 | 0 | 0 | 0 | 0 | 0 | 0 | 0 | 0 |
| **CRF11_cpx** | 4 | 0.3 | 1 | 0 | 0 | 0 | 0 | 0 | 1 | 2 | 1 | 0 | 0 | 3 | 0 | 0 | 0 | 0 | 0 | 0 | 0 |
| **CRF18_cpx** | 1 | 0.1 | 1 | 0 | 0 | 0 | 0 | 0 | 1 | 0 | 0 | 0 | 0 | 0 | 0 | 0 | 0 | 0 | 0 | 0 | 0 |
| **CRF19_cpx** | 1 | 0.1 | 0 | 0 | 0 | 0 | 0 | 0 | 0 | 0 | 1 | 0 | 0 | 1 | 0 | 0 | 0 | 0 | 0 | 0 | 0 |
| **CRF43_02G** | 236 | 16.4 | 33 | 11 | 1 | 0 | 0 | 0 | 45 | 138 | 34 | 0 | 0 | 172 | 8 | 0 | 8 | 3 | 0 | 0 | 19 |
| **CRF49_cpx** | 1 | 0.1 | 1 | 0 | 0 | 0 | 0 | 0 | 1 | 0 | 0 | 0 | 0 | 0 | 0 | 0 | 0 | 0 | 0 | 0 | 0 |
| **D** | 9 | 0.6 | 2 | 0 | 0 | 0 | 0 | 0 | 2 | 4 | 2 | 0 | 0 | 6 | 0 | 0 | 1 | 0 | 0 | 0 | 1 |
| **G** | 119 | 8.3 | 24 | 6 | 0 | 0 | 0 | 0 | 30 | 55 | 25 | 1 | 0 | 81 | 4 | 1 | 2 | 0 | 0 | 1 | 8 |
| **URF** | 328 | 22.7 | 78 | 13 | 0 | 0 | 2 | 1 | 94 | 163 | 46 | 0 | 3 | 212 | 9 | 0 | 11 | 0 | 2 | 0 | 22 |
| **Total** | **1442** | **100** | **346** | **64** | **1** | **1** | **22** | **1** | **435** | **697** | **216** | **1** | **15** | **929** | **34** | **1** | **35** | **5** | **2** | **1** | **78** |

N, number of sequences; LAG, Lagos; IBA, Ibadan; OND, Ondu; OSU, Osun; ENU, Enugu; ABV, Abuja; NIG, Niger; NC, Other North central areas; KAD, Kaduna; YOB, Yobe; MAI, Maiduguri; KAN, Kanu; ADA, Adamawa; BOR, Borno

### Subtype determination

The Nigerian *pol* sequences were aligned with the 2010 All M group (A-K + Recombinants) reference sequence dataset (http://www.hiv.lanl.gov/) using the Clustal algorithm, followed by manual editing in MEGA6^27,28^. The HIV-1 subtype/CRF assignment was determined with maximum-likelihood (ML) phylogenetic analysis in Garli v0.98^29^, applying the General Time Reversible (GTR) substitution model to infer the ML phylogeny. Branch support was estimated using the approximate likelihood ratio test with the Shimodaira-Hasegawa-like procedure (aLRT-SH) as implemented in the PhyML 3.0^30^. Branches with aLRT-SH support >0.90 were considered statistically supported^25^.

### Recombination analysis

Putative unique recombinant forms (URFs) and sequences that were difficult to type were analysed by Bootscan in Simplot^31^. Briefly, *pol* sequences were aligned with the LANL 2010 HIV-1 reference subtypes for G, CRF4302G and CRF02AG (parental sequences). Recombination breakpoints were identified using a sliding window size of 300 bp and step size of 50 bp.

In order to define the structure and distribution of the breakpoints across the alignment, we plotted a line graph of the relative frequency of the breakpoints. The K-means univariate-clustering algorithm as implemented in the ‘Ckmeans.1d.dp’ R package was used to define hotspots for recombination^32^. The gap statistic method implemented in the ‘factoextra’ R-package^33^was employed to estimate the groups based on similarity in breakpoint positions obtained from the Simplot analysis. The recombination hotspot positions were then used to identify groups of three or more URFs with one or more similar breakpoint positions. Finally, for the URFs to be defined as potential new CRFs, we performed a maximum likelihood phylogenetic analysis to assess the epidemiological relatedness among the sequences^34^.

### Cluster analysis

A previously described BLAST approach was used to construct a reference sequence dataset for each subtype/CRF, separately^25,35^. Briefly, we initially constructed a reference set containing at least eight non-Nigerian sequences from the BLAST search of each sequence belonging to the different CRF/subtype group. Redundant sequences from each reference dataset were removed using an in-house Perl script and the Emboss 6.6.0.0 package skip redundancy^36^. Nigerian transmission clusters were defined as clusters with aLRT-SH support >0.90 and ≥80% Nigerian sequences^25,37^. Clusters of two sequences were defined as dyads, 3-14 sequences as networks, and >14 sequences as large clusters^38^.

### Dating and Phylogeographic analysis

The temporal signal in each dataset were assessed by TempEst using the transition-transversion versus divergence plots^39^. To determine informative substitution rate priors for analysis of the main transmission clusters in Nigeria, we randomly sampled 150 sequences for each subtype/CRF, respectively. The evolutionary rates were estimated in BEAST v1.8^40^using the SRD06 model^41^with a relaxed uncorrelated lognormal clock model^42,43^. Markov chain Monte Carlo (MCMC) simulations were run for 30×10^7^chain steps, subsampling parameters every 1000 steps. Convergence was assessed in Tracer.v.1.6 (Effective Sample Sizes (ESS) ≥100)^44^. Estimated subtype/CRF-specific evolutionary rates were then used as priors in the subsequent analyses.

We used a Bayesian discrete phylogeographic approachwith a MCMC length of 300 million steps in BEAST v1.8, sampling every 30,000^th^step, to reconstruct the spatial dynamics of HIV-1 for the large clusters identified^40,45^. BEAGLE was used to improve run time of likelihood calculations, and Tracer v1.6 was used to assess convergence of the runs (ESS ≥100)^44,46,47^. The demographic history of the viral population, past growth rates, and effective population sizes were inferred using the Bayesian Skygrid model^48^. Priors for the TreeModel Root Height were estimated using the Bayesian Skyline model^49^. If applicable, detailed growth rates were estimated using the exponential growth rate model.

Symmetric and asymmetric continuous time Markov chain models were used to model the location exchange process and the parsimonious description of the location exchange rates was inferred using the Bayesian stochastic search variable selection (BSSVS) procedure^24,49,50^. A robust counting approach as implemented in BEAST was used to estimate the forward and reverse viral movement events between locations along the branches of the posterior tree distributions^51^. Well-supported movements were summarized using SPREAD v1.0.7 based on a Bayes factor cut-off >3^52-54^. The percentage of viral movements was summarized using R^55^.

All files and scripts are available from the authors upon request.

### Statistics

Linear by linear association test (LBL) was used to analyze trends over time, using IBM SPSS V22.0 Armonk, NY: IBM Corporation.

### Ethics

Approvals were obtained from the local and national institutional review boards affiliated to the treatment sites and the international collaborating sites including the University of Maryland, Baltimore, Harvard University, University of Amsterdam and US center for diseases control.

## RESULTS

### CRF02_AG, CRF43_02G and subtype G were the major circulating strains in Nigeria

We analyzed 1442 HIV-1 *pol* sequences collected from four geopolitical zones in Nigeria between 1999 and 2013 (Table 1). The phylogenetic analysis showed that the CRF02_AG was the most common strain (44% of the analyzed sequences), followed by CRF43_02G (16%), subtype G (8%) and CRF06_cpx (4%). A large proportion of the sequences (23%) were unique recombinant forms (URFs), whereas the remaining sequences were minor variants (each variant accounting for <2%).

Most sequences were from Abuja (697 sequences, 48%), followed by Lagos (346 sequences, 23%) and Jos (216 sequences, 15%). The distribution of different subtypes/CRFs varied within these regions/states with fewer CRF02_AG infections following a North East direction from Lagos (Figure 1). Analysis of trends over time showed an overall decrease in the proportion of CRF02_AG infections in Nigeria (from 55% in 2006 to 38% in 2013, p=0.015, LBL, Figure 2). Moreover, the analysis also showed an increase in the proportion of unique recombinant forms (URFs) from 16% to 32%, 2005-2009 (p=0.015, LBL).

**Figure 1.**
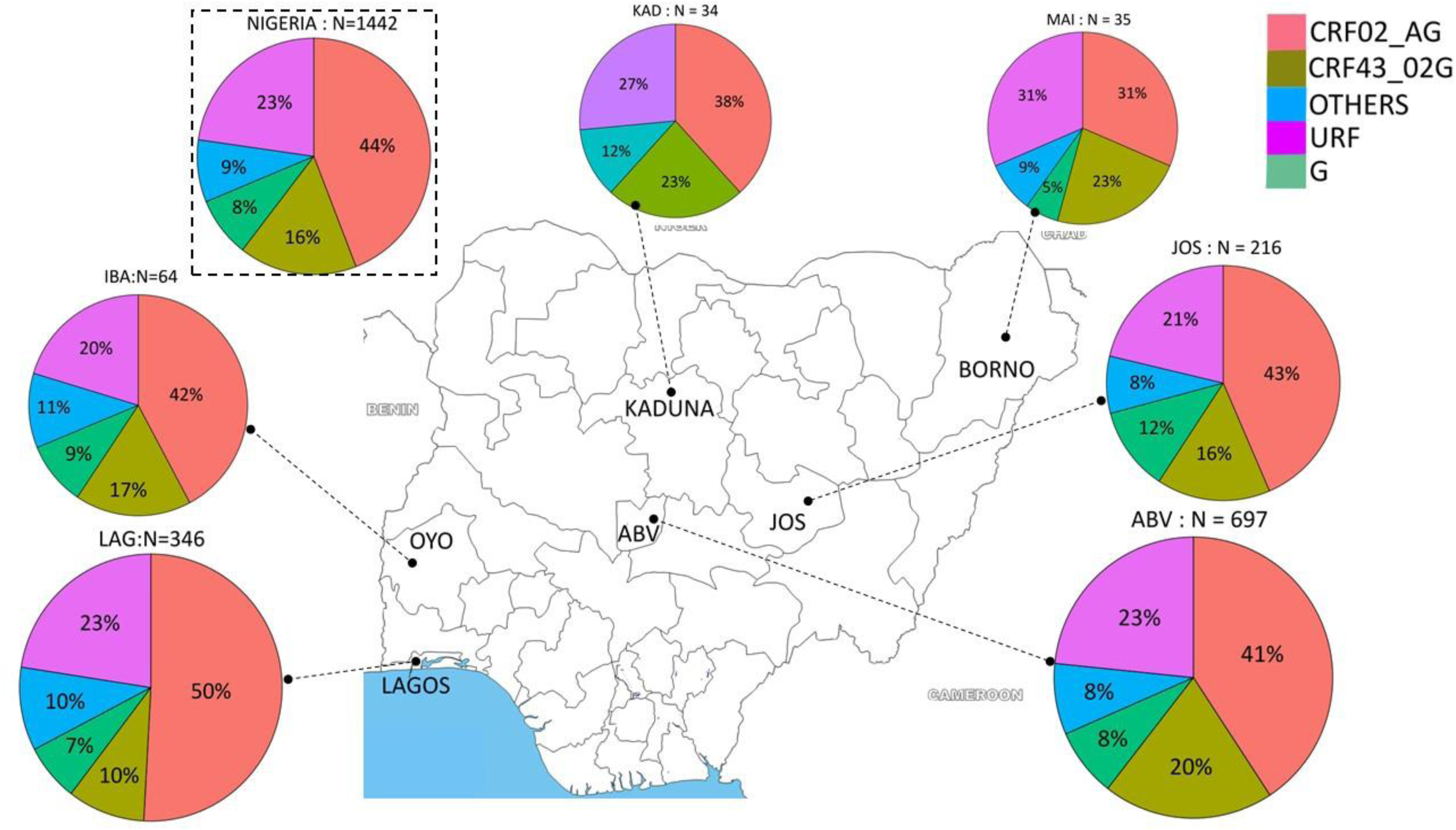
Prevalence of subtypes/CRFs among HIV-1 infected individuals collected from different locations in Nigeria. Molecular diversity of HIV-1 in different zones/regions in Nigeria. The location of the sampled regions are shown by the black dot with a line leading to diversity pie-chart for the city in that particular region. The colors in the pie-chart are defined in the key. Abbreviations for the towns: ABV, Abuja; KAD, Kaduna; MAI, Maiduguri; LAG, Lagos; IBA, Ibadan (Based on the map from https://d-maps.com/).

**Figure 2.**
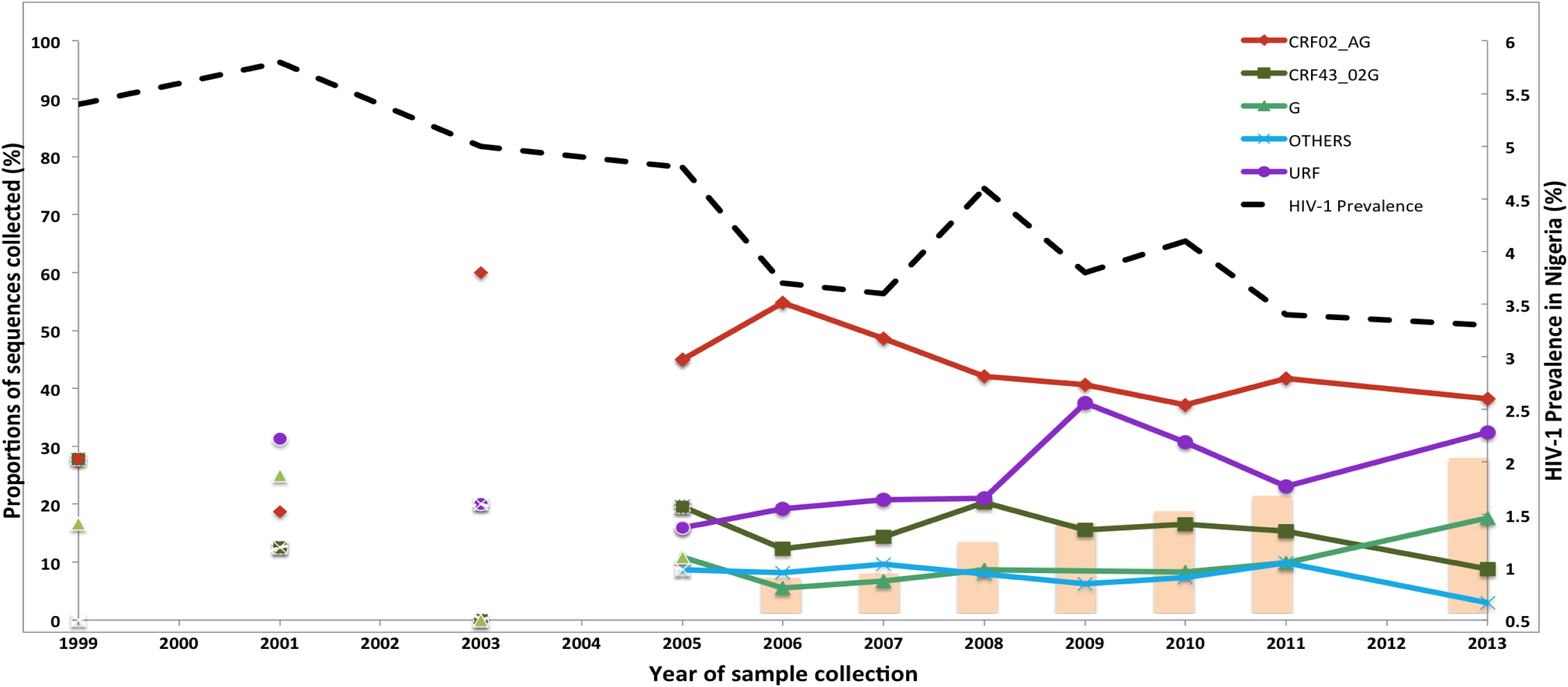
Dynamics of subtype/ CRF proportions, ART coverage and overall country HIV-1 prevalence over time. The subtype/ CRF proportion displayed as overall percentage of sequences collected from Nigeria per year. Few sequences (< 20 per year) were collected from 1999-2004 and their proportions had no effect on the LBL association tests. No sequences were collected in 2012. The orange bars represent the proportion of HIV-1 infected individuals that were receiving ART in the respective years. The change in proportion of different subtypes/CRFs over time was not significant for subtype G (p = 0.711, LBL), CRF43_02G (p = 0.497, LBL), CRF02_AG (p = 0.323, LBL) and CRF06_cpx (p = 0.015, LBL). However the increase of URFs over time was significant (p = 0.015, LBL). The y-axis shows the proportions (%) of sequences collected over time for a particular subtype/CRF or the proportion (%) of patients receiving ART. The x-axis represents the time period in years from 1999-2013. The z-axis shows the HIV-1 prevalence in Nigeria over time.

### Four potential recombination hotspots in the *pol* region

A detailed recombination analysis was performed on 210 of the 310 sequences classified as URFs. These sequences were initially selected from the maximum likelihood phylogenetic trees if they branched off close to the root between two and had long branches. There were 655 breakpoint positions recorded among the 210 sequences, with some sequences having more than one breakpoint. These positions were plotted as a frequency plot of breakpoints along the alignment to identify hotspots for recombination (Supplementary Figure 1). Alignment positions around 285-315 (HXB2 K03455 positions: 2538-2568), 503-534 (2756-2787), 720-775 (2973-3028) and 923-956 (3176-3209) were identified as potential recombination hotspots. To define these positions more precisely, we used the IQR of the optimal univariate K-median clustering algorithm with the number of K clusters determined by the gap-statistic method^56^(Supplementary Figure 2): Recombination hotspot I: 294-312 (HXB2 K03455 positions: 2547-2565); II: 503-533 (2756-2786); III: 729-805 (2982-3058); and IV: 931-957 (3184-3210) (Supplementary Table 1). One-hundred-and-thirty-nine of the 210 (66%) sequences had a recombination breakpoint in the hotspot I region; 104/210 (50%) in hotspot II region; 58/210 (28%) in hotspot III region; and 59/210 (28%) in hotspot IV region. The hotspot positions were unique independent recombination events from the original breakpoint positions as previously identified for the parental sequences of CRF43_02G (HXB2 K03455 positions: 1266, 3325, and 6097) and CRF02_AG (HXB2 K03455 positions: 2391, 3275, and 4175). We identified seven groups of URFs with 3-10 sequences with similar breakpoint positions (Supplementary Figure 6) and that were not epidemiologically linked.

### Cluster analysis

To determine transmission clusters of the major circulating strains within Nigeria, we analyzed the three dominating forms CRF02_AG, CRF43_02G, and subtype G, respectively. In total, 206 subtype G sequences were analyzed (119 Nigerian and 87 non-Nigerian). The phylogenetic analysis showed four dyads, one network, and one large Nigerian subtype G cluster (consisting of 81 Nigerian and 11 non-Nigerian sequences, Table 2 and Supplementary Figure 3). Analyses of the 1161 CRF02_AG sequences (636 Nigerian and 555 non-Nigerian, Supplementary Figure 4) resulted in 12 dyads, 12 networks and six clusters (Table 2). Finally, we analyzed 295 CRF43_02G sequences (236 Nigerian and 59 non-Nigerian, Supplementary Figure 5) and all the Nigerian sequences clustered monophyletically (SH-aLRT=0.99). The majority of the reference sequences obtained by the BLAST approach did not cluster within the CRF43_02G cluster, suggesting that CRF43_02G mainly circulates in Nigeria.

**Table 2.** Number of clusters in the different subtype / CRF groups.

| **Subtype / CRF** | **Dyads<sup>a</sup>** | **Networks<sup>b</sup>** | **Large clusters<sup>c</sup>** | **Total** |
| --- | --- | --- | --- | --- |
| G | 8 (4) | 8 (2) | 94 (1) | 104 |
| CRF02_AG | 24 (12) | 76 (12) | 336 (6) | 439 |
| CRF43_02G | 0 (0) | 0 (0) | 295 (1) | 295 |
| **Total** | **32** | **84** | **725** | **841** |

<sup>a</sup>Dyads: Clusters of 2 sequences
<sup>b</sup>Networks: Clusters of 3-14 sequences
<sup>c</sup>Large clusters: Clusters with more than 14 sequences
The brackets ( ) indicates the number of clusters observed for the different groups.

### Dating the origin of five main Nigerian transmission clusters

To further dissect the Nigerian HIV-1 epidemic, we focused on the identified large Nigerian clusters; one subtype G, three CRF02_AG and one CRF43_02G clusters. Analysis of the temporal signal showed that all clusters had a correlation coefficient above 0.3, indicating a positive correlation between genetic distance and sampling time, and thus retained sufficient phylogenetic signal to conduct coalescence analyses^57^.

The median time to most recent common ancestor (tMRCA) of the Nigerian subtype G cluster was estimated to 1979 (95% HPD: 1967-1980); the CRF02_AG clusters to 1961 (95% HPD: 1946-1974), 1968 (95% HPD: 1958-1978) and 1960 (95% HPD: 1946-1973) for cluster 1, 2 and 3, respectively; and the CRF43_02G cluster to 1980 (95% HPD: 1975-1983) (Figure 3). The median HIV-1 evolutionary rates were estimated to 2.1×10^−3^substitutions/site/year (s/s/y) (95% HPD: 1.7-2.5×10^−3^s/s/y) for the Nigerian subtype G cluster; 1.4×10^−3^s/s/y (95% HPD:1.1-1.9× 10^−3^s/s/y), 1.3×10^−3^s/s/y (95% HPD: 1.4-1.8×10^−3^s/s/y), 1.2× 10^−3^s/s/y (95% HPD: 0.9-1.6×10^−3^s/s/y) for the CRF02_AG cluster 1,2 and 3, respectively; and 4.2×10^−3^s/s/y (95% HPD:3.5-4.8×10^−3^s/s/y) for the CRF43_02G cluster (Figure 3).

**Figure 3.**
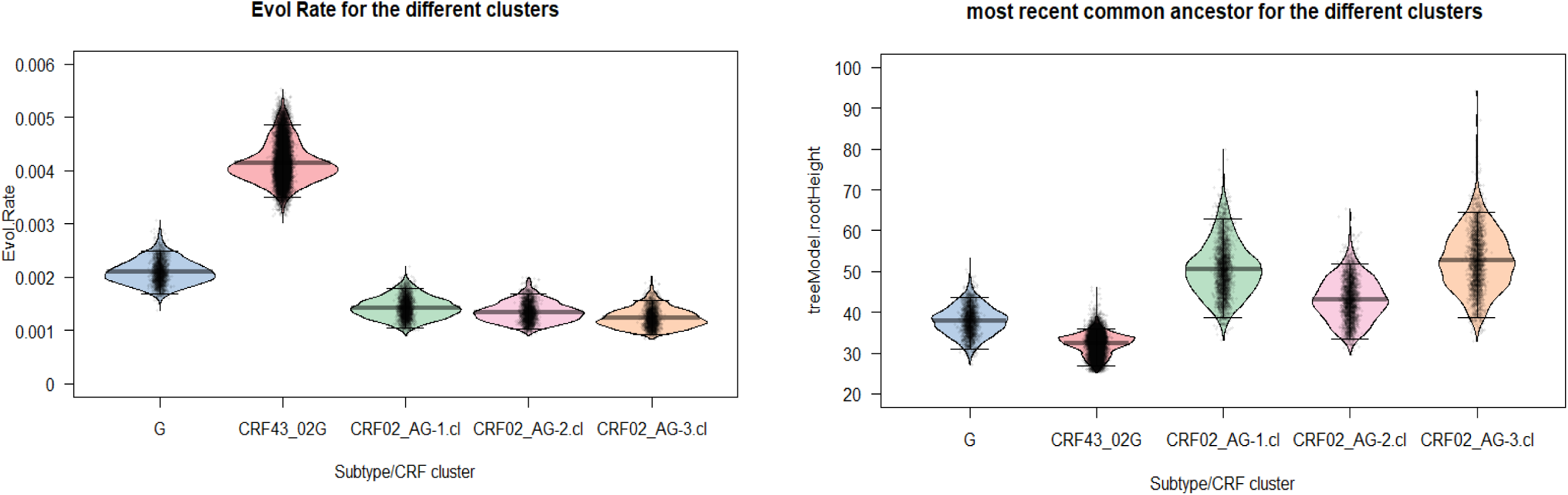
Demographic history for the different Nigerian clusters. Pirate plots for the evolutionary rate and time to the most recent common ancestor since 2013 for the different 5 clusters in Nigeria. Each bean in the plot has mean indicated as a bold horizontal black line and the Bayesian 95% highest density interval as thin horizontal lines. The raw data from the sampling interval during Bayesian analysis is indicated by black dots inside the beans. The CRF43_02G had a slightly higher evolutionary rate compared to the other clusters.

To control for bias in the phylogeographic analysis due to oversampling of some locations in the dataset in relation to what is reflected in the epidemic, we randomly selected sequences from different regions based on their HIV-1 prevalence and population growth over time in the different geographic regions. In these analyses, the median tMRCA of the Nigerian subtype G cluster was estimated to 1987 (95% HPD: 1982-1992); and the CRF02_AG clusters to (95% HPD: 1960-1983), 1972 (95% HPD: 1973-1981), 1961 (95% HPD: 1952-1979) for cluster 1, 2 and 3, respectively.

### Disentangling the demographic history of five main Nigerian transmission clusters

The Bayesian Skygrid analysis indicated that the number of effective infections (i.e. the number of individuals contributing to new HIV-1 infections over time^58^) in the Nigerian subtype G epidemic underwent a fast exponential growth between the 1970s and the mid-1990s with an increase to 10,000 effective infections, followed by a marginal decrease with minor fluctuations from the mid-1990s (Figure 4). The median growth rate was 30% per year (95% HPD: 18%-42%). The three clusters representing the CRF02_AG epidemic displayed a similar pattern with a slow increase in growth rate from the 1980s to 2000s. The median CRF02_AG growth rates were estimated to 22% per year (95% HPD: 13%- 32%) for cluster 1, 18% per year (95% HPD: 10%-26%) for cluster 2, and 24% (95% HPD: 11%-38%) for cluster 3. Finally, the CRF43_02G cluster also showed an increase in effective HIV-1 infections from 1980 to 2000 (from 100 to 10,000 effective HIV-1 infections), followed by a relatively sharp decrease from mid-2000 and forward (Figure 4). The median growth rate was 30% per year (95% HPD: 18%-42%).

**Figure 4.**
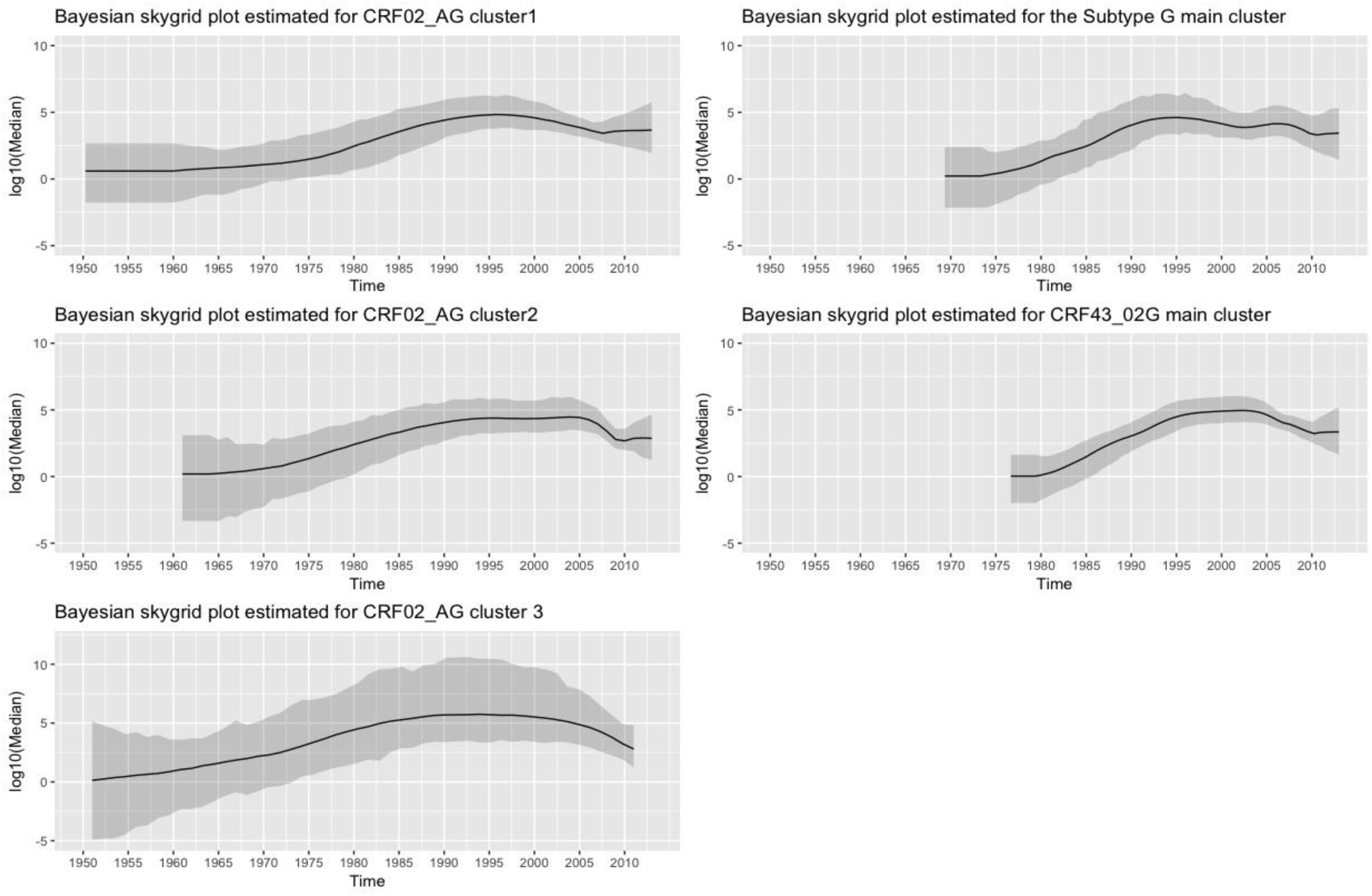
Skygrid plots for the different clusters. Phylodynamic analyses of HIV-1 *pol* gene for subtype G, CRF02_AG, CRF43_02G clusters isolated in Nigeria. Bayesian skygrid plots representing the changes in the effective population size of the virus (*N_e_*)(log) over time. The solid lines represent the estimated median log effective population size and the gray shaded areas represents the 95% HPD intervals.

### Phylogeographic dispersal of HIV-1 in five main Nigerian transmission clusters

Next, we sought to investigate the spatio-temporal process of the HIV-1 spread in Nigeria using symmetric and asymmetric continuous time markov chain (CTMC) phylogeographic models with the BSSVS procedure in BEAST. The two models gave similar estimations. Results from the asymmetric analysis for subtype G showed a high statistical support for epidemiological linkage between Kaduna and other regions: Abuja (Bayes Factor [BF]=11302), Ibadan (BF=6), Yobe (BF=1609), and Jos (BF=11302). For CRF02_AG clusters, Abuja was connected to Ibadan, Jos, Kaduna, Lagos, Oyo and Maiduguri (all BFs >4). Based on the posterior distribution of the location of origin, the most probable origin of Subtype G, CRF43_02G and CRF02_AG was Abuja (Posterior probability: 0.98). However, these results should be interpreted with caution due to the low number of sequences from some locations.

Since the dataset contained unbalanced distribution of samples in terms of location, we conducted a control analysis to assess the robustness of our results to over-represenations of samples from particular regions in Nigeria (Table 1). We selected samples based on population size and HIV-1 prevalence and run the CTMC analysis on the subsample. This analysis indicated that the most probable root state for all the strains where outside Nigeria.

### Rates of viral lineage migration

The rates of viral lineage migration within Nigeria were estimated using a robust counting approach to infer the history of viral movements and epidemiological links between different locations and the locations that contributed most to the dispersal of HIV-1 subtypes within Nigeria. For subtype G, HIV-1 dispersal from Abuja was estimated at 42% (95% HPD: 35-48%) and the highest viral migrations from Abuja were to Jos (17%, 95% HPD: 12-22%), outside Nigeria (10%, 95% HPD: 8-13%), and Lagos (7%, 95% HPD: 4-10%). Similar results were found for the CRF02_AG clusters, except for slightly higher migration rates from Abuja to Lagos for cluster 1 (30%, 95% HPD: 25-34%, Figure 5). The CRF02_AG dispersal from Abuja was estimated to 61%, 52% and 21% from cluster 1, 2 and 3, respectively.

**Figure 5.**
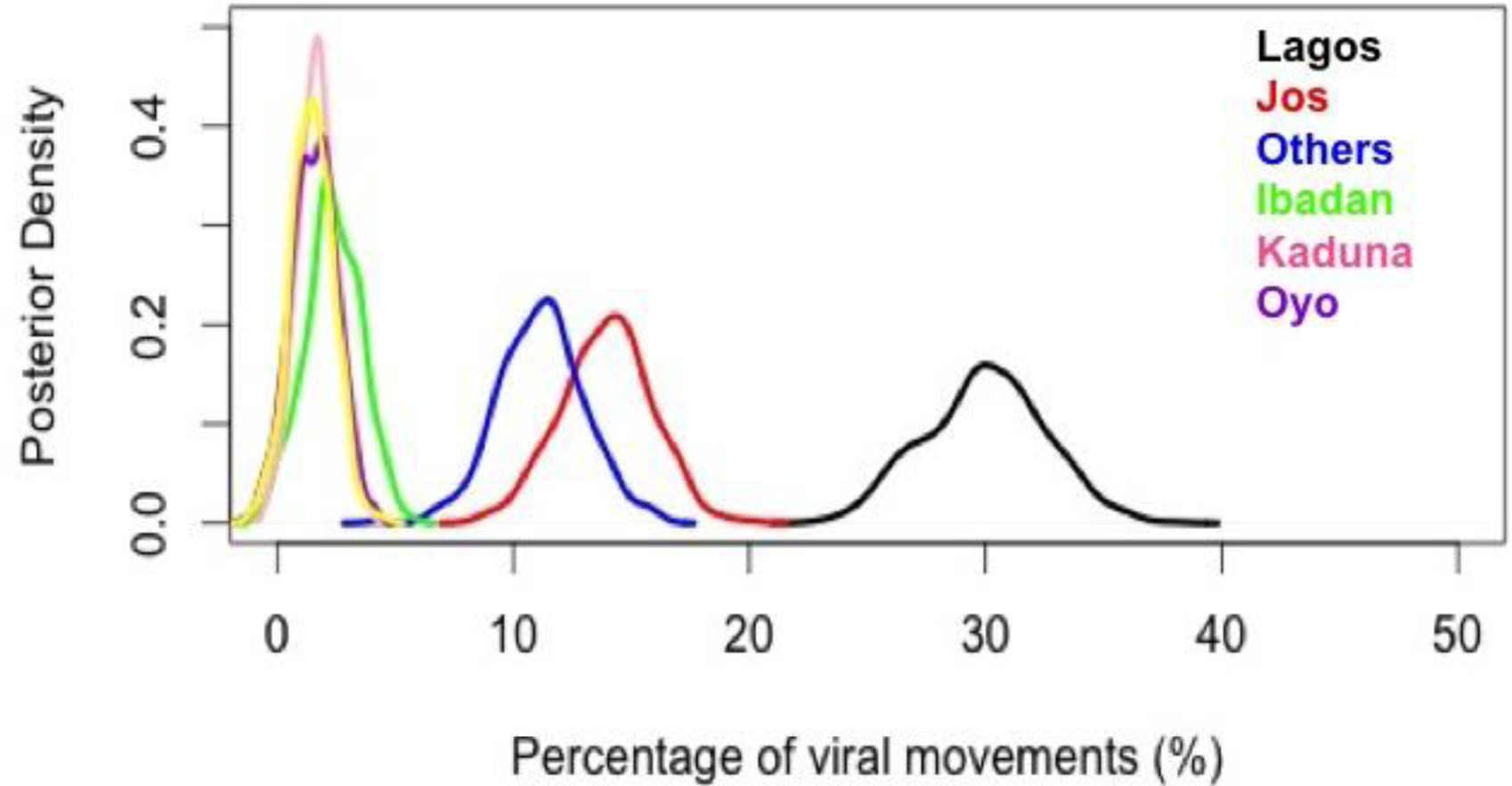
Estimated percentages of CRF02_AG migration events from Abuja to each location in the NG, obtained using cluster 1. The density plot for the viral movements from the Abuja (most probable root location) to Lagos, Jos, Kaduna and other towns. Lagos and Jos had the highest percentage of viral movements from Abuja for the CRF02_AG Cluster 1. Infections from outside Nigeria accounted for 10% of the viral movement.

## DISCUSSION

In this study, we aimed to provide a better understanding of the history and spread of the HIV-1 subtypes/CRFs circulating in Nigeria. We applied state of the art phylogenetic approaches on a large set of HIV-1 sequences from individuals of the five most populated geopolitical zones in Nigeria to characterize the Nigerian HIV-1 epidemic and to obtain new insights about its origin and disseminations patterns. In line with previous studies, we identified CRF02_AG, CRF43_02G and subtype G as the most prevalent strains (previous estimates ranging from 19-60% for CRF02_AG^8,9,11-13^and 22-50% for subtype G^9-11,14,15^). The prevalence of CRF43_02G has only been reported in one previous study from Abuja (estimated to 19%)^10^. These large variations and discrepancies are likely explained by analysis of local geographic regions, and that different subtyping tools differ in accuracy in assigning the correct subtype/CRF (CRFs are often particularly challenging)^59^. It could also be the result of a sampling not reflecting the real prevalence regions and/or over periods.

For both subtype G and CRF43_02G, we found one well-supported monophyletic clade, respectively. Each clusters harbored the vast majority of the Nigerian sequences from those strains suggesting introductions by a single strain or a limited number of strains that grew out to dominate the epidemic. In contrast, three large Nigerian clusters were found for the CRF02_AG strain indicating multiple introductions that grew out to separate sub-epidemics.

We observed one large transmission cluster with 68% of all Nigerian subtype G sequences and a coalescent estimate on the time of emergence at 1979 (1967-1980). This estimate is consistent with that subtype G cluster (GWAII) described by Delatorre *et al.* in 2014 and it is approximately six years before the first AIDS cases was identified Nigeria^60^. This indicates that HIV-1 had spread in Nigeria already before the first AIDS case was identified in the country. The three independent transmission clusters for CRF02_AG that originated from within Nigeria had coalescent estimates on the time of emergence from 1960-1980. Our analysis indicated that the one of the CRF02_AG cluster had an earlier origin than the estimated tMRCA for subtype G (i.e. the putative parental strain). This supports a previous study that CRF02_AG is in fact the parental strain of subtype G^61^. Interestingly, the CRF43_02G was first fully described and isolated in Saudi Arabia in 2008^62^. Our analysis indicated that it emerged in Abuja already in the early 1980’s. In addition, the majority of the CRF43_02G strains in Genbank was of Nigerian origin. Considering the large prevalence of both the putative parental strains CRF02_AG and subtype G, it is therefore plausible that CRF43_02G first emerged in Nigeria. The molecular clock estimates on the date of introduction, from the control analysis accounting for the sampling bias, corresponded well with all the five clusters. This implies that the sampling location had limited effect on these estimations.

The demographic analysis indicated an increase in number of effective infections 1970-1995 in all five clusters. This is in line with a rapid population growth during the same period in Nigeria (from 56 to 108 million people, www.worldometers.info/world-population/nigeria-population). Moreover, this increase was followed by a decline in number of effective infections which seem to coincide with when free ART was introduced in 2006^6^, and when Nigeria registered a sharp decrease in HIV-1 prevalence (from approximately 6% to 3%)^3,6^. However, despite this decrease in HIV-1 prevalence, our analysis indicated a relatively high number of co-circulating strains. Our analysis also indicated that urban areas like Abuja and Lagos might represent major hubs of HIV-1 transmissions.

The recombination analysis indicated several URFs and potentially novel CRFs. Inter-subtype/CRF recombination does not occur randomly on the HIV-1 genome as its frequency varies over the genome with several so-called hotspots for recombination^63,64^. One main hotspot for recombination in the *pol* region (found in 105 sequences) was close to the *pol PR-RT* border (positions 2547-2565 in the HIV-1 HXB2 reference genome, K03455). This hotspot has previously been reported in a study on HIV-1 subtype B^65^. The hotspot positions II and IV has also been reported elsewhere^66^. The seven identified groups of URFs represents potential novel CRFs circulating in Nigeria or second generation recombinants with different recombination patterns. However, this needs to be confirmed by full-length sequencing. To date, 18 distinct subtype G related CRFs have been described and published worldwide indicating the diversity in Subtype G sequences (http://www.hiv.lanl.gov/).

This is the first study to dissecting the Nigerian molecular epidemiology with a country-wide set of HIV-1 sequences. Understanding the underlying processes and factors that influence HIV-1 transmission, demographic history and migration patterns are pivotal to understand the epidemic potential of different HIV-1 strains. Ultimately, this may also inform population-based surveillance of infectious diseases.

## NOTES

### Acknowledgments

We thank all the study participants and the collaborating health centers in Nigeria.

## Author contributions

J.N, N.N and J.E. interpreted the data and were responsible for the overall study design. N.N and J.E. were responsible for the overall project coordination. H.R., B.C., P.D, A.A, M.C were medically and organisationally responsible for the clinical sites and collected key epidemiological data of the study participants. J.N, N.F. and J.E analysed the data and contributed in statistical analyses. J.N. and J.E. wrote the manuscript. All authors read and approved the manuscript.

## SUPPLEMENTARY DATA

### Characterisation of the HIV-1 Molecular Epidemiology in Nigeria: Origin, Diversity, Demography and Geographic Spread

Jamirah Nazziwa^a^, Nuno Faria^b^, Beth Chaplin^c^, Holly Rawizza^c^, Patrick Dakum^d^, Alash’le Abimiku^d^, Man Charurat^d^, Nicaise Ndembi^d^and Joakim Esbjörnsson^a^

Department of Laboratory Medicine, Lund University, Lund, Sweden^a^

Department of Zoology, University of Oxford, Oxford, United Kingdom^b^

Department of Immunology and Infectious disease, Harvard T.H School of Public Health, Boston, USA^c^

Institute of Human Virology, Abuja, Nigeria^d^

### Files in this Data Supplement

Supplementary Table 1. Summary of the median, mean and interquartile ranges for the 4 hotspot regions in the pol alignment using the univariate K-means clustering Algorithm

Supplementary Figure 1. Potential recombination breakpoint hotspots detected in the pol alignment

Supplementary Figure 2. Optimal cluster determined by the gap statistic method

Supplementary Figure 3. Maximum likelihood phylogenetic tree for Subtype G sequences

Supplementary Figure 4. Maximum likelihood phylogenetic tree for CRF02_AG sequences

Supplementary Figure 5. Maximum likelihood phylogenetic tree for CRF43_02G sequences

Supplementary Figure 6. Groups of sequences with similar recombination breakpoint

Legends for supplementary figures

### SUPPLEMENTARY TABLES

**Supplementary Table 1.** Summary of the median, mean and interquartile ranges for the four hotspot regions in the *pol* alignment using the univariate K-means clustering Algorithm.

|  | I | II | III | IV |
| --- | --- | --- | --- | --- |
| N | 221 | 203 | 115 | 116 |
| Min. | 147.00 | 415.00 | 656.00 | 864.00 |
| 1st Qu. | 294.00 | 503.00 | 729.00 | 931.00 |
| Median | 305.00 | 522.00 | 768.00 | 948.00 |
| Mean | 306.38 | 519.96 | 760.44 | 941.24 |
| 3rd Qu. | 312.00 | 533.00 | 805.00 | 957.25 |
| Max. | 398.00 | 633.00 | 858.00 | 980.00 |

### SUPPLEMENTARY FIGURE LEGENDS

**Supplementary Figure 1.**
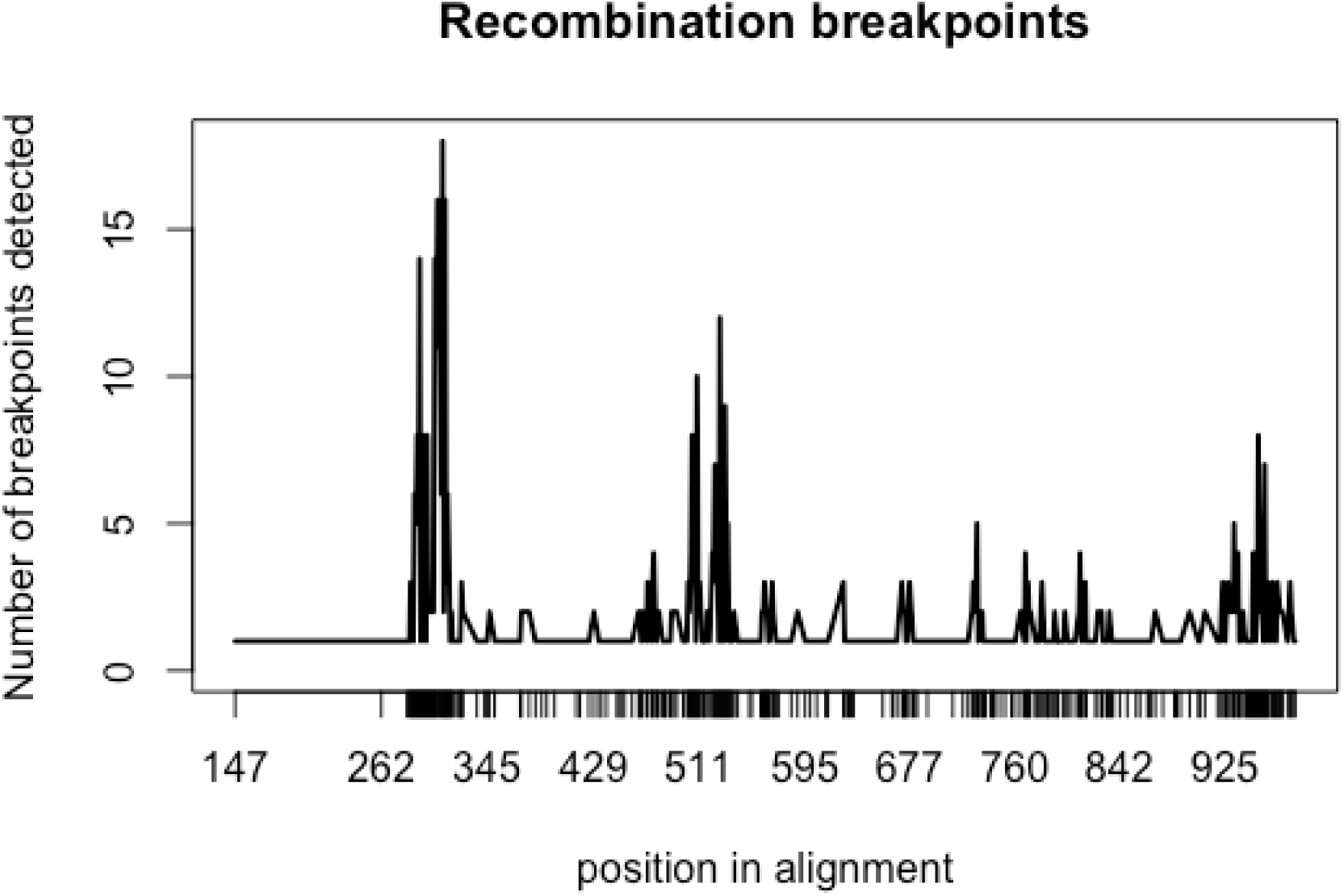
Potential recombination breakpoint hotspots detected in the *pol* alignment. Small vertical lines at the bottom of the graph indicate breakpoint positions along the alignment. Breakpoints detected in each window of 300 nucleotides along the alignment were counted and plotted (solid line).

**Supplementary Figure 2.**
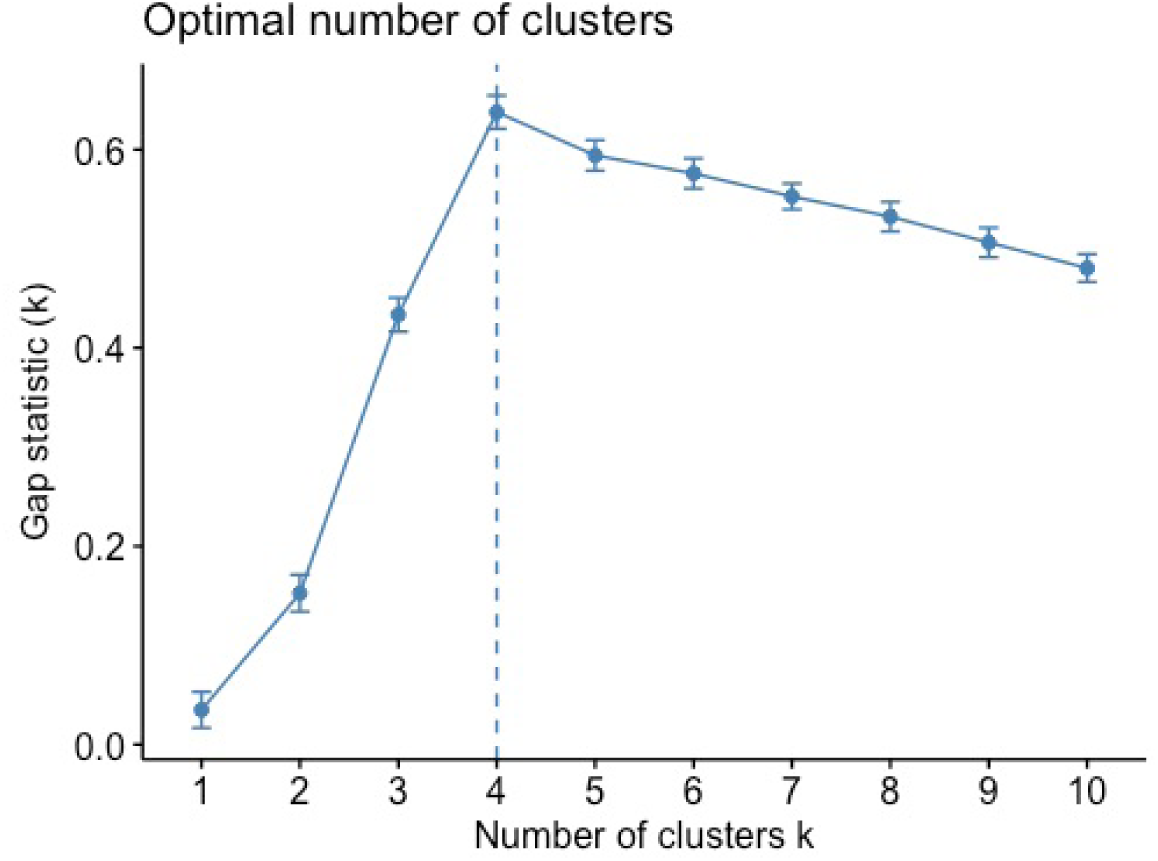
Optimal cluster determined by the gap statistic method. The gap statistic compares the total intra-cluster variation for different values of k with their expected values under null reference distribution of the data. This plot provides the gap statistic and standard error, identifying k=4 as the optimal cluster with the highest gap statistic.

**Supplementary Figure 3.**
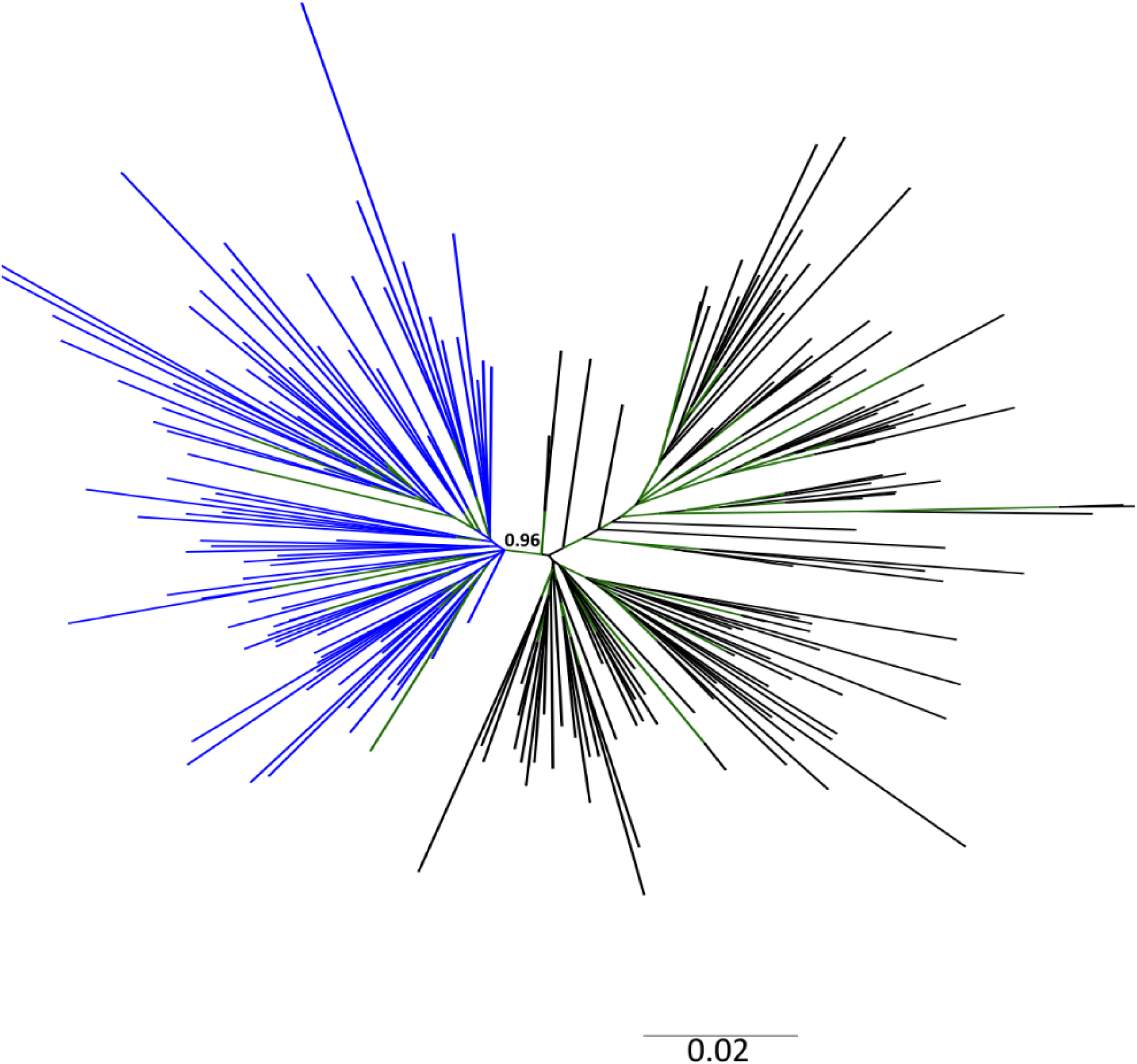
Maximum likelihood phylogenetic tree of the pol region for Subtype G sequences (n = 203). The blue braches indicate Nigerian sequences while the green indicate an SH-aLRT branch support above 0.9. The black branches indicate the non-Nigerian sequences. We observed one large NG cluster (in blue) that was considered for further analysis

**Supplementary Figure 4.**
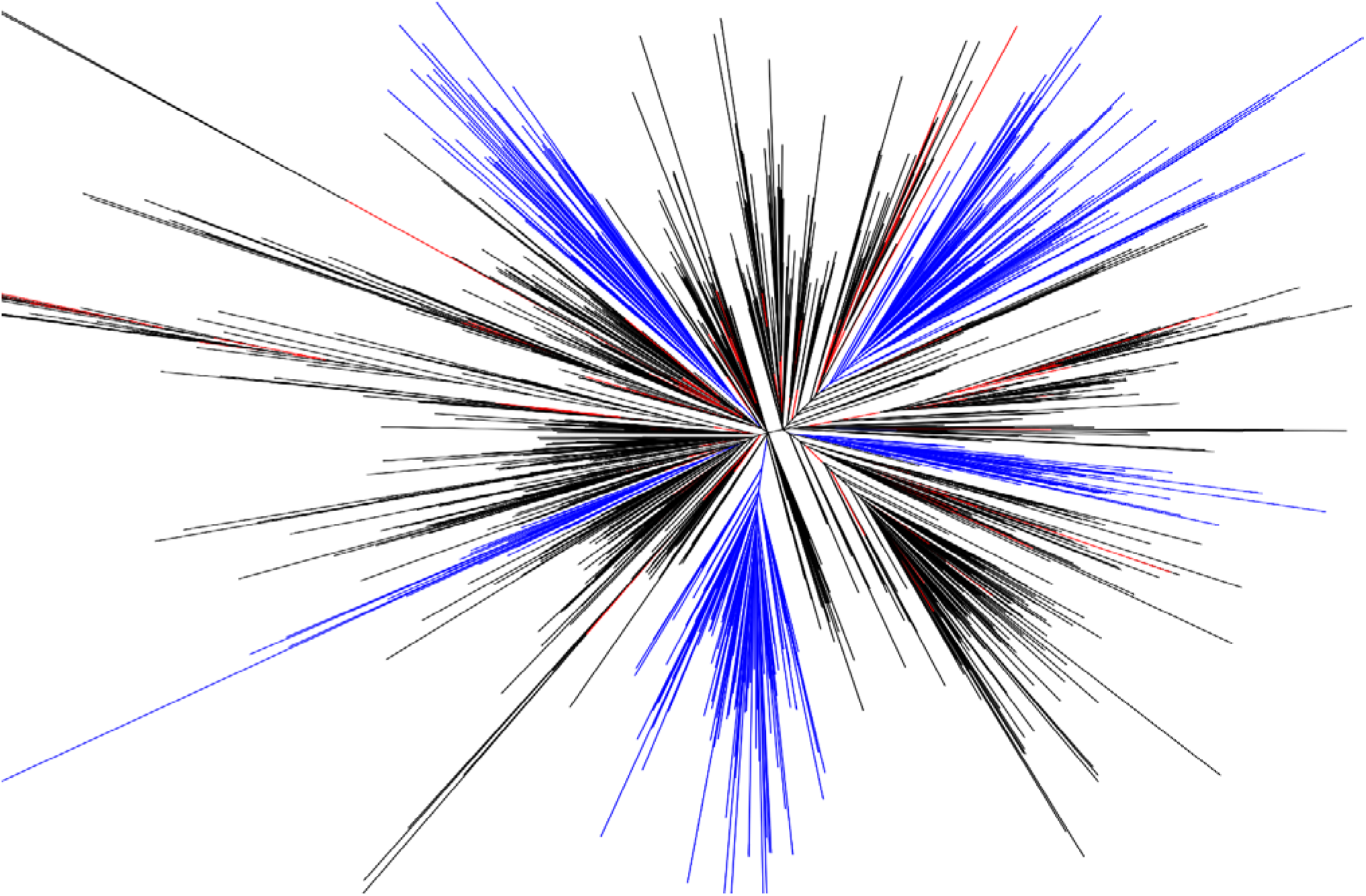
Maximum likelihood phylogenetic tree for CRF43_02G sequences (n = 667). The blue braches indicate Nigerian sequences while the green indicate an SH-aLRT branch support above 0.9. The black branches indicate the non-Nigerian sequences. We observed one large NG cluster (in blue) that was considered for further analysis. The purple sequences were later classified as URFs

**Supplementary Figure 5.**
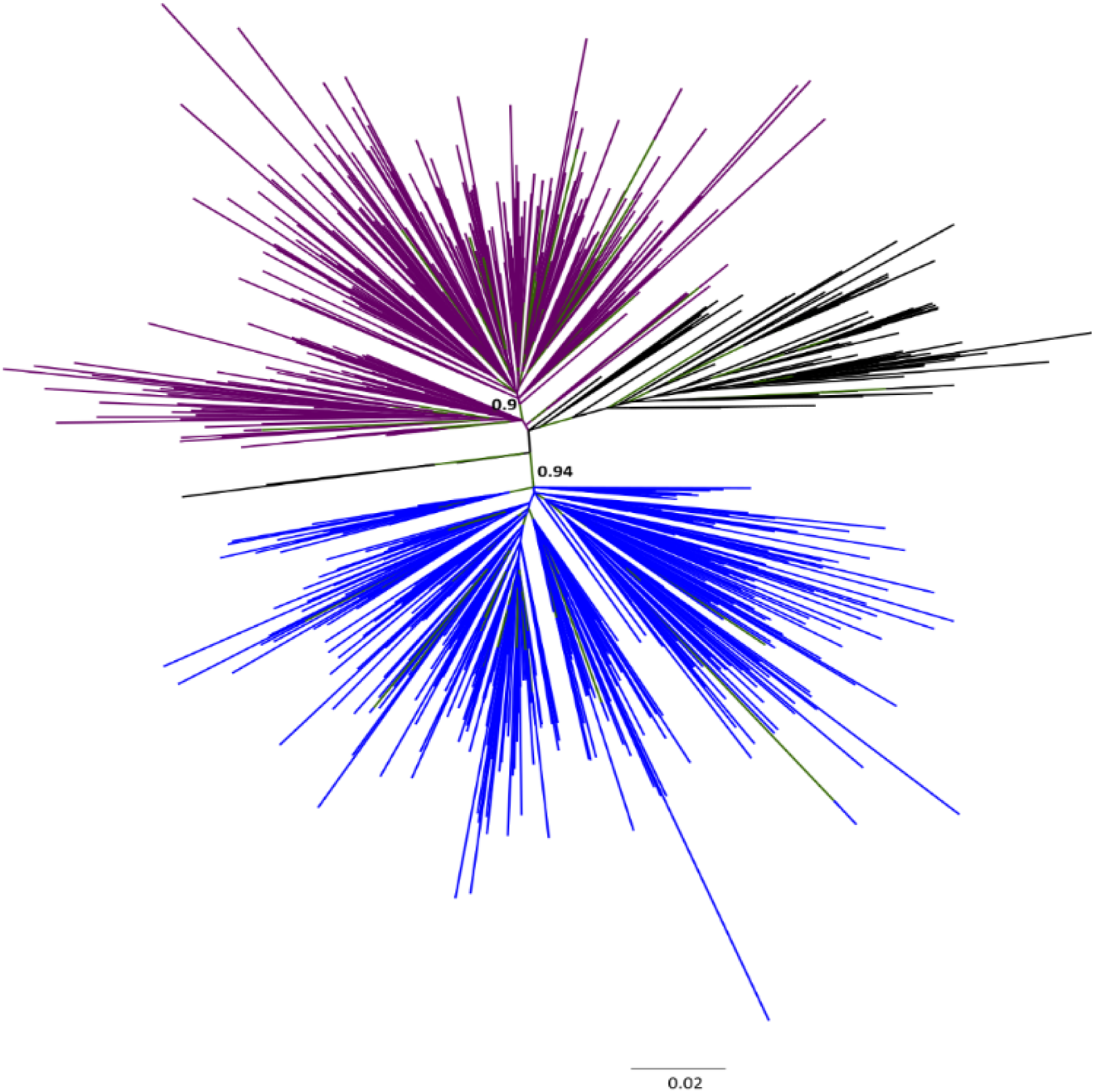
Maximum likelihood phylogenetic tree for CRF02_AG sequences (n = 1170). The blue braches indicate Nigerian sequences while the green indicate an SH-aLRT branch support above 0.9. The black branches indicate the non-Nigerian sequences. We observed 3 large Nigerian clusters (in blue) that were considered for further analysis.

**Supplementary Figure 6. Groups of sequences with similar recombination breakpoint patterns that could be potential new CRFs.**

